# Improvisation and Live Accompaniment Increase Motor Response and Reward During a Music Playing Task

**DOI:** 10.1101/2023.09.28.559982

**Authors:** Anna Palumbo, Karleigh Groves, Eva Luna Muñoz Vidal, Alan Turry, Robert Codio, Preeti Raghavan, Heidi Schambra, Gerald T. Voelbel, Pablo Ripollés

## Abstract

Music provides an abstract reward that can enhance learning and motivation in humans. While music is often combined with exercise to improve performance and to upregulate mood, the relationship between music-induced reward and motor output is poorly understood. Here, we study music reward and motor output at the same time by capitalizing on music playing. Specifically, we investigate the effects of music improvisation and live accompaniment on motor, autonomic, and affective responses. Thirty adults performed a rhythm tapping task while (i) improvising or maintaining the beat and (ii) with live or recorded accompaniment. Motor response was characterized by acceleration of hand movements (accelerometry), wrist flexor and extensor muscle activation (electromyography), and the number of beats played. Autonomic arousal was measured by tonic response of electrodermal activity (EDA) and heart rate (HR). Affective responses were measured by a 12-item Likert scale. The combination of improvisation and live accompaniment, as compared to all other conditions, significantly increased acceleration of hand movements and muscle activation, as well as participant reports of enjoyment during music-playing. Improvisation, regardless of type of accompaniment, increased the number of beats played and autonomic arousal (including tonic EDA responses and several measures of HR), as well as participant reports of challenge. Importantly, increased motor response was associated with increased enjoyment during music improvisation only and not while participants were maintaining the beat. The increased motor responses achieved with improvisation and live accompaniment have important implications for enhancing dose of movement during music-based interventions for stroke rehabilitation.

**Significance Statement:** Music provides a rewarding stimulus and improves motor performance and learning. However, the relationship between music reward and motor output is poorly understood. Here, we show that music improvisation with live accompaniment increased acceleration and muscle activation during movement, and that this increase in motor response was associated with increased enjoyment only when improvising. These findings are important for developing music interventions that target improved motor performance and learning in exercise and physical rehabilitation.

## Introduction

Reward plays an important role in motor performance and learning (Wulf and Lewthwaite, 2016). Monetary reward during motor tasks increases force production, reduces reaction time, improves recall of motor skills, increases neural activation in reward-related areas and modulates corticospinal excitability (Pessiglione et al., 2007; Widmer et al., 2016; Bundt et al., 2019). Dopaminergic signaling between reward and motor brain areas optimizes motor skill learning in non-human animals (Molina-Luna et al., 2009; Hosp et al., 2011). In everyday life, *music* provides an abstract reward that is frequently combined with exercise and athletic training to improve mood and performance (Laukka and Quick, 2013; Hallett and Lamont, 2017). Listening to music improves affect, endurance, speed, power output, movement kinematics, and cardiovascular efficiency (Simpson and Karageorghis, 2006; Karageorghis et al., 2009; Terry et al., 2012; Belkhir et al., 2019; Stork et al., 2019; Terry et al., 2020; Bonassi et al., 2023). Music listening activates reward-related brain areas, and musical pleasure is modulated by dopamine release (Blood and Zatorre, 2001; Menon and Levitin, 2005; Salimpoor et al., 2011; Ferreri et al., 2019). Musical groove also activates reward and motor-related brain areas (Matthews et al., 2020). Music playing (i.e., using musical instruments) has been utilized in motor rehabilitation to improve recovery following acquired brain injury, leading to increased motor function, reduced depression and improved positive affect, and increased activation in the affected sensorimotor cortex (Grau-Sanchez et al., 2013; Raghavan et al., 2016; Ripolles et al., 2016; Magee et al., 2017; Palumbo et al., 2022). Critically, music playing supports higher levels of motor recovery among individuals with greater sensitivity to musical reward (Grau-Sanchez et al., 2018). All this evidence suggests that increasing reward during music playing improves motor outcomes. However, the mechanisms supporting these effects remain unclear. Here we capitalize on music playing to study the putative link between movement and reward. We examine two avenues for enhancing reward and motor output during music playing: music improvisation and live accompaniment.

Music improvisation involves creating or adapting music in the moment, providing choice about what to play (Turry, 2009; Loui, 2018). Choice enhances motor performance and learning, even when the choice is unrelated to the task, and increases motivation and activation in reward-related brain areas (Wulf et al., 2014; Lewthwaite et al., 2015; Murayama et al., 2015; Chua et al., 2018; An et al., 2020). In this vein, choice in combining sounds increases enjoyment and interest (Baur et al., 2018). Music improvisation, compared to playing a pre-defined set of notes, increases activation in the sensorimotor cortex and in brain regions related to executive functions (Beaty, 2015; de Aquino et al., 2019; Vuust et al., 2022). Creativity in music improvisation correlates with grey matter in the hippocampus (Arkin et al., 2017). These findings indicate the potential for music improvisation to influence cognition (e.g., attention, learning), motor responses, and reward during music-playing.

Live accompaniment creates a social experience during music playing. Positive social interactions contribute to reward (Krach et al., 2010), and cooperative learning of motor tasks increase intrinsic motivation, positive affect, self-efficacy, and motor learning as compared to competitive or non-interactive learning (Kaefer and Chiviacowsky, 2022). As compared to recorded music, live music also modulates heart rate and mood, including self-reports of reduced anxiety and improved vigor, (Bailey, 1983; Bush et al., 2021; but see Belfi et al., 2021). Together, these findings indicate that live music may reduce feelings of anxiety and their associated physiological responses and that social interactions during live music playing may improve reward and motor learning.

Here, we compare the effects of improvisation and live accompaniment in a music-playing task in healthy participants. We predict that improvisation combined with live accompaniment will produce the highest levels of motor activity, positive affect, and autonomic arousal. We also predict that increased motor responses will correlate with increased self-reports of reward-related affective responses during music-playing, thus supporting the proposed link between reward and motor performance.

## Methods

### Participants

Thirty healthy adults were recruited from the New York University community, ages 18-58 (M = 22.9, SD = 8.11; 19 women). Sample size was based on previous studies with 13-21 participants demonstrating that rhythmic improvisation increases the number of notes played and the activation of motor planning and executive brain areas (Berkowitz and Ansari, 2008; de Aquino et al., 2019). All participants were either enrolled in university (n = 24) or had already earned an undergraduate or graduate degree and were currently employed (n = 4). Participants self-reported their race as Asian (n = 14), White/Caucasian (n=8), Black/African American (n=2), or Mixed (n=2), with 3 participants not reporting their race. Twenty seven percent (n=8) of participants reported Hispanic ethnicity. Inclusion criteria included right-handedness with no history of brain injury. Exclusion criteria included musical anhedonia, as indicated by a score of < 65 on the Barcelona Music Reward Questionnaire (Mas-Herrero et al., 2013), or current employment as a professional musician. The participants gave written informed consent as per the institutional review board at New York University and were paid for their participation.

### Music Playing Task

Participants played a drum along with piano accompaniment for four one-minute songs, written specifically for this study. Each of the four songs were piano compositions based on one of the following musical styles: March, Waltz, Salsa, and Rock. Prior to playing, there was a familiarization period. During this period, participants first listened to a recording of each song and then practiced playing along. While listening, participants were instructed to remain as still as possible. While practicing, participants were asked to play along with the recorded piano accompaniment and find the beat, or pulse, of the music that felt most natural to them. If participants were not able to identify what a “beat” or “pulse” was in the music, they were given several examples of where the beat can fall in music, and then continued practicing until they identified a beat. Practicing continued until participants identified a steady beat that they preferred to play, which followed the beat of the music, and which they felt able to maintain. After the familiarization period, participants were instructed to play the drum along with the piano accompaniment by following one of two *music playing conditions*: (1) Beat or (2) Improvise. During the *Beat* condition, participants were instructed to maintain a steady rhythm that they preferred to play and which they felt followed the beat, or pulse, of the music. They were not required to play the same beat as they identified in the familiarization period, and they were not required to choose the same beat each time they played. However, each time they selected a beat to play during the Beat condition, participants were asked not to change the rhythm they played as much as possible throughout the song. In contrast, during the *Improvise* condition, participants were invited to vary their rhythm freely, as desired for personal expression throughout the song. In this way, the Beat and Improvise conditions both represented personal preference, but only the Beat condition was constrained to stay the same throughout the song. Participants played the drum with piano music provided with one of two *accompaniment conditions* (1) Recorded or (2) Live. During the R*ecorded* condition, accompaniment was played from a recording of the song. The same accompaniment recordings were used for all participants. During the *Live* condition, accompaniment was performed live by the same music therapist who played in the recordings. The live accompaniment was provided in a different manner based on the music playing condition. During the Beat condition, the live accompaniment was played following the piano composition as closely as possible. During the Improvise condition, the music therapist played the piano composition responsively, to reflect the improvised rhythmic patterns played by the participant. This responsive playing followed guidelines used in clinical music therapy to support participant’s sense of being heard and supported in their music making (Nordoff et al., 2007). Importantly, responsive playing was required to stay within the overall structure of the composition, such that the melodic and harmonic structures retained the contour of the original composition. The music therapist remained in the room in all conditions, regardless of whether a recording was being played. Piano and drums were played on Musical Instrument Digital Interface (MIDI) devices, including the Roland SPD-One Percussion MIDI drum pad, tuned to the sound of snare drum, and a Roland FP-60x digital keyboard. Both MIDI inputs were recorded to individual tracks in Logic Pro, with real time sound output to a Roland KC-400 150 watt 12-inch keyboard amp.

### Measures

Baseline measures evaluated demographic characteristics of participants and musical reward-sensitivity and training. Outcome measures included both subjective behavioral self-reports provided immediately following music playing and continuous objective physiological and motor measures collected during music playing. The behavioral measures evaluated affective and physical responses to music-playing. The continuous measures evaluated motor and autonomic responses. Motor responses were measured using electromyography (EMG), accelerometry, and the number of beats played. Autonomic responses were measured using electrodermal activity (EDA) and electrocardiogram (ECG). EMG, accelerometry, EDA, and ECG data were collected using a MP160 data acquisition module (BIOPAC Systems Inc.) and AcqKnowledge software (see additional details below). All signal processing for this data was completed in MATLAB R2022b. Custom code for processing of physiological data in MATLAB is available upon request to the first author.

### Baseline Characteristics

Baseline screening included demographic information - including age, sex, race, ethnicity, and education level. Baseline screening also measured factors known to play a role in response to music-playing tasks, including sensitivity to musical reward and musical training. Sensitivity to music reward was measured via the Barcelona Music Reward Questionnaire (BMRQ), a reliable five-factor scale (Mas-Herrero et al., 2013). Music training was measured via the music training subscale of the Goldsmith Music Sophistication Index (Gold-MSI), a self-report scale with strong internal consistency (Cronbach’s alpha = 0.903;Müllensiefen et al., 2014).

### Behavioral Correlates of the Music Playing Experience

After each music-playing task, participants provided responses for twelve questions using a 5-point Likert scale. These questions were related to their affective and physical response, as well as their perception of the music (Table 1). Principal component analysis (PCA) was used to identify the main components of participant responses across these questions. The number of main components were determined using parallel analysis. Participant loadings on these main components, rather than responses to each Likert scale question, were analyzed in relation to participant experience of conditions (see Results).

**Table 1:**
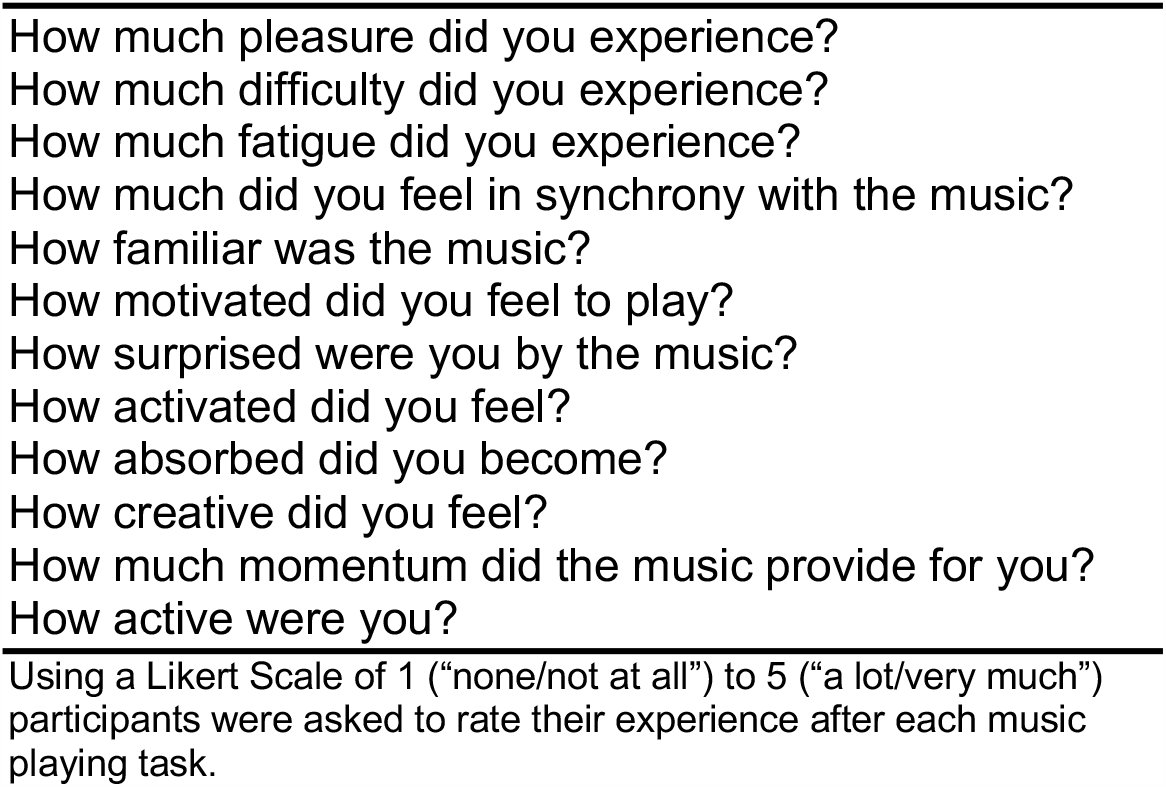
Experience of Music Survey Items.

### Electromyography

EMG was measured via surface electrodes placed over the wrist flexor carpi radialis and extensor carpi radialis longus on the arm used to play the drum. Prior to electrode placement, the skin was cleaned with alcohol. EMG signals were filtered with a 10-400 Hz band-pass filter, constructed by convolving a 2^nd^ order high pass Butterworth filter and 3^rd^ order low pass Butterworth filter (Aluru et al., 2014). The signals were then rectified by taking the root mean square of 10ms windows of the signal. Signals were normalized by dividing by the participant’s maximum voluntary contraction (MVC; Aluru et al., 2014). The MVC was measured by providing resistance to movement at about 45° of rotation through wrist extension and flexion, during which participants were asked to exert the maximum possible force against the resistance.

MVC signals were filtered and rectified as described above, and the final MVC value was calculated as the average amplitude of the maximum one-second region of the signal.

To ensure background noise did not influence the overall EMG average, movement-related peaks in normalized EMG amplitude were identified by alignment with peaks in the Z component of the accelerometry signal (see below) using custom code. The lowest 10% of peaks in EMG amplitude were dropped to eliminate noise, and plots were visually inspected to ensure proper alignment between EMG and accelerometry signals. The average of movement-related peaks in EMG amplitude was calculated. Since participants played the drum with slightly different arm positions, the results present findings for the average wrist flexor and extensor activation, rather than considering each muscle separately. EMG signals were recorded at a sampling rate of 2000 samples/second.

### Accelerometry

A triaxial accelerometer was secured to the back of the hand playing the drum using a Velcro strap, with placement under the metacarpophalangeal joints and between the third and fourth fingers. Accelerometry signals were acquired at a sampling rate of 2000 samples/second, except for the first 14 participants, for whom the y component was sampled at 250 samples/second due to a technical problem. To correct for this, all other accelerometry signals were down sampled to 250 Hz. A 4^th^-order Butterworth high pass filter with a cut-off at 0.5 Hz was then used to remove the gravitational component (Hurd et al., 2013). Leading and trailing samples were dropped due to filter and resampling artifacts, with an overall sample loss of less than 10% of the signal. The x, y, and z components of the resulting accelerometer signal were then transformed into magnitude of acceleration taking the square root of the sum of the squares of each component. The total acceleration was calculated as the sum of the magnitude of acceleration over the entire signal (Hurd et al., 2013).

### Number of Beats Played

Number of beats played, (i.e., taps on the drum) during each music condition was calculated from the MIDI recordings. MIDI files were loaded into MATLAB, and the MIDI toolbox 1.1 was used to calculate the number of beats played (Eerola and Toiviainen, 2004).

### Electrodermal activity

EDA electrodes were placed on the index and middle finger of the hand not playing the drum, with the hand resting on a table. We instructed participants to move the hand with the EDA electrodes as little as possible to avoid motion artifacts. Tonic response of EDA was calculated as the mean of the signal for each trial (Braithwaite et al., 2013). Music-playing responses were normalized for each participant by subtracting the tonic response to the listen-only condition from each trial. Five participants were dropped from the EDA analysis due to hyperhidrosis that caused ceiling effects in the EDA signal (n = 4) or insufficient signal (n = 1).

### Electrocardiography

ECG was recorded using three electrodes placed on the torso. Raw ECG signals were detrended, and heartbeats were identified as the R wave maxima of the QRS complexes. Noise was removed using custom code, and ECG signals were visually inspected to ensure that QRS complexes were accurately identified. Three participants were dropped from ECG analyses due to insufficient signal-to-noise ratio. Beats per minute (BPM) were calculated as the sampling rate (in samples/sec) divided by the average samples per beat, multiplied by 60 (sec/min; Gorr, 2023). Heart rate variability (HRV) was calculated as the root mean square of successive differences (RMSSD) between R wave maxima (Shaffer et al., 2016; Shaffer and Ginsberg, 2017). ECG signals were recorded at 2000 samples/second.

### Experimental Design and Statistical Analysis

Using a crossover design, all participants completed all four combinations of music playing and accompaniment conditions (Figure 1) with each of the four songs, for a total of 16 music playing trials. Participants were randomized to one of four condition orders (Figure 1.1 in Extended Data). The same condition order was repeated for each of the four songs played by each participant, and all conditions were completed for one song before moving to the next song. For each outcome, linear mixed models were calculated with fixed effects including the music playing condition (Improvise vs. Beat), the music accompaniment condition (Live vs. Recorded), and their interaction, with control variables for musical training, sensitivity to music reward, and song, and a random intercept for subject (Bates et al., 2015). For EDA and ECG, total acceleration was additionally included in the model as a control, to eliminate motion as an explanation for increased autonomic response. Post-hoc comparisons of significant effects were adjusted for multiple tests using the Tukey method (Lenth et al., 2019).

**Figure 1:**
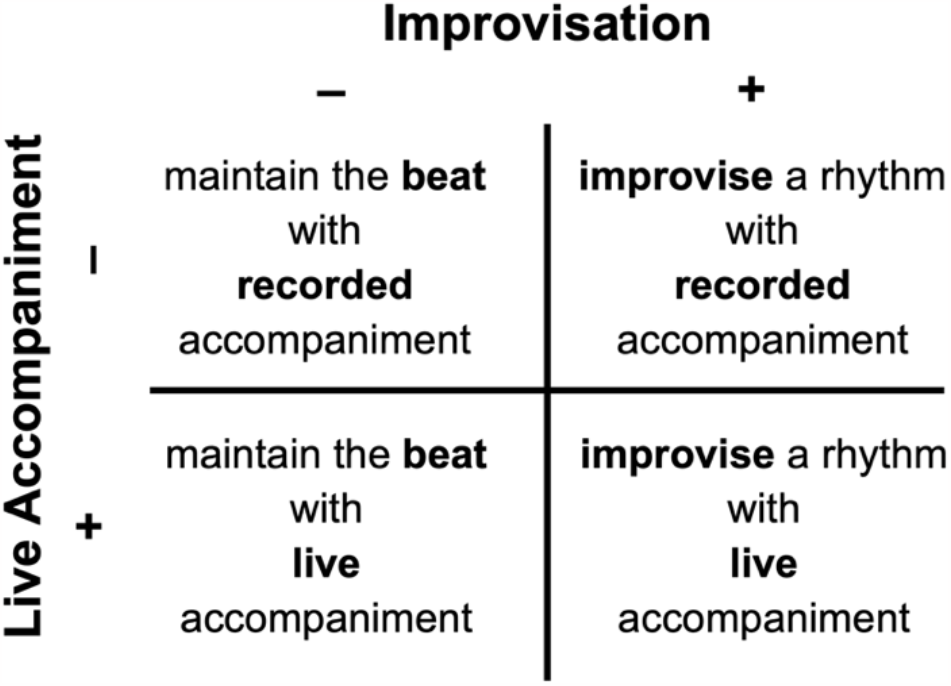
Experimental Design. Using a 2x2 design, each participant completed all four combinations of the music playing tasks, including with or without improvisation (music playing conditions: Improvise vs. Beat) and with or without live accompaniment (accompaniment conditions: Live vs. Recorded).

To evaluate the association between reward-related behavioral measures and motor responses (accelerometry, EMG, and beats played), we again used linear mixed models (Bates et al., 2015). As a measure of reward, we used the first principal component of participant responses to the 12-item survey related to the music playing experience. This principal component includes positive loadings for pleasure, as well as other responses related to enjoyment (see Results), and therefore more fully accounts for participant responses related to reward as compared to the single rating of pleasure. Fixed predictors included motor response (we computed 3 different models for our 3 different measures: accelerometry, EMG, and number of beats played), music playing condition (Improvise vs. Beat), music accompaniment condition (Live vs. Recorded), and their three-way interaction, as well as control variables for musical training, music reward, and song, with a random intercept for subject. The three-way interaction in this model accounts for potential modulation of the association between motor responses and enjoyment by the four music tasks. Post-hoc comparisons of linear trends were evaluated by averaging over other factor level predictors in the model (Lenth et al., 2019). To account for multiple comparisons, significance levels with and without Bonferroni correction are reported for the association between Enjoyment and the three motor responses.

For all analyses of linear mixed models, Wald Chi Square tests were used to evaluate significance of main effects and interactions. All analyses were done in R (packages: psych, factoextra, lme4, foreign, aod, car, emmeans, emtrends, ggplot2, ggeffects).

## Results

### Participant Musical Background

Participant scores on the BMRQ (M = 84.93, SD = 9.08) were in the range of the general population (Mas-Herrero et al., 2013; M = 78.42, SD = 10.47, n = 857). Similarly, participant scores on the Gold-MSI musical training subscale (M = 25.40, SD = 10.82) were in the range of the general population (Müllensiefen et al., 2014; M = 26.52, SD = 11.44, n = 147,633). These findings indicate our sample approximates the general population in relationship to sensitivity to musical reward and musical training.

### Behavioral Correlates of the Music Playing Experience

PCA of responses revealed two main components of music playing experience (Figure 2). The first component (PC1) accounted for 38.05% of the variance in the survey responses and was termed “Enjoyment” to summarize positive loadings for participant experiences of feeling pleasure, momentum, activated, motivated, creative, active, and absorbed. The second component (PC2) accounted for 18.82% of the variance in the survey responses and was termed “Challenge” to summarize positive loadings of participant experiences of difficulty, fatigue, and surprise, and negative loadings of participant perception of the music as familiar and in synchrony with their playing.

**Figure 2:**
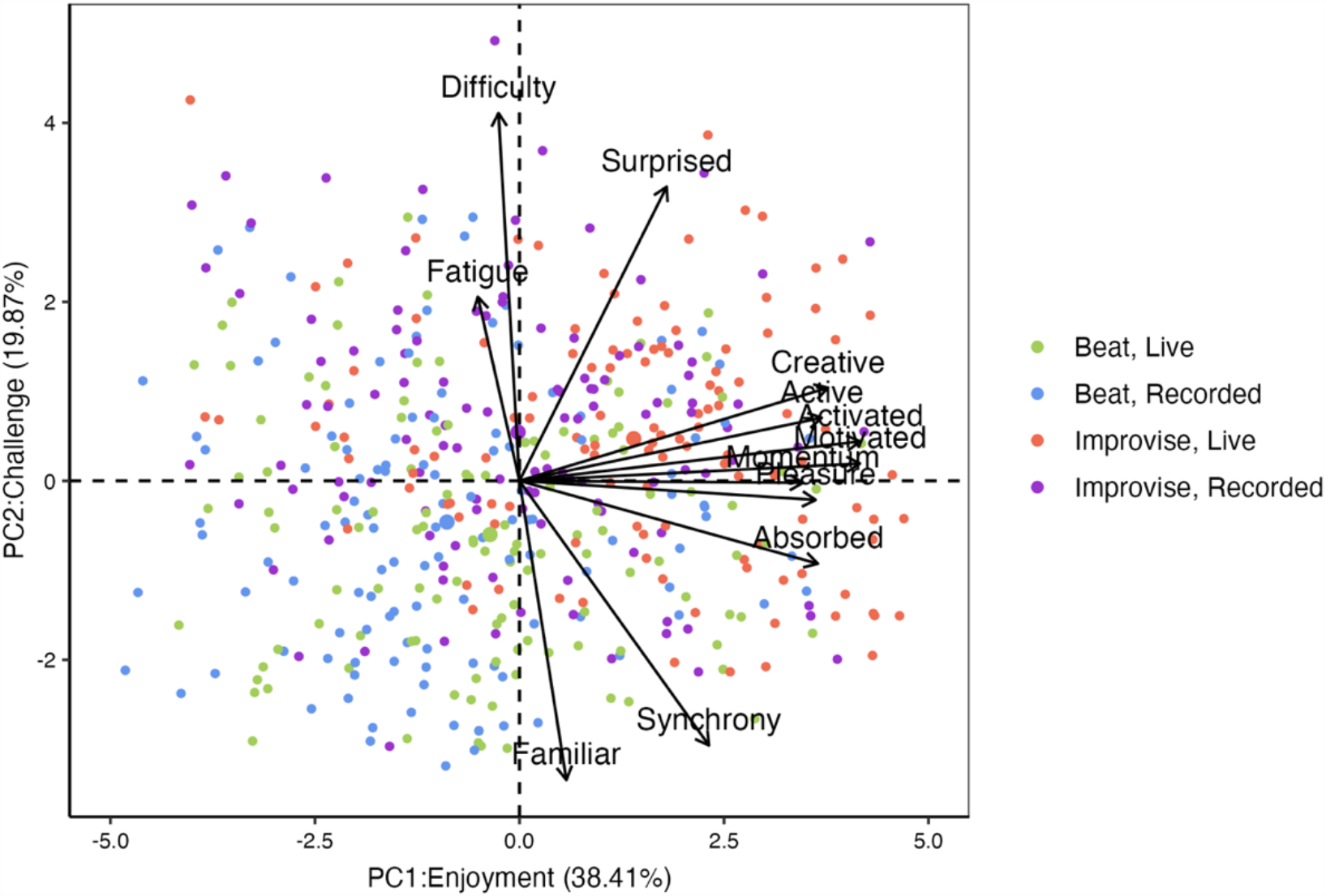
Principal Component Analysis of Music Experience Survey. Biplot of principal components derived from the 12-item music experience survey. The first principal component includes responses related to Enjoyment, accounting for 38.41% of the variance in survey responses. The second principal component includes responses related to Challenge, and accounted for 19.87% of the variance in participant responses.

For Enjoyment (see Figure 3), there were significant main effects of music playing condition (Improvise vs. Beat; χ^2^ (1) = 15.18, p = 0.0001) and accompaniment condition (Live vs. Recorded; χ^2^ (1) = 7.73, p = 0.005), as well as for song (χ^2^ (3) = 33.92, p < 0.00001). In addition, greater sensitivity to musical reward was related to greater Enjoyment (χ^2^ (1) = 4.51, p = 0.034). There was also a significant interaction between playing condition and accompaniment condition on Enjoyment (χ^2^ (1) = 14.53, p = 0.0001). Pairwise comparisons for song, corrected for multiple tests, revealed less Enjoyment for the March as compared to the Rock (t(444) = -5.33, p < 0.0001) and Salsa (t(444) = -3.20, p = 0.0006) compositions, but not the Waltz (t(444) = -1.57, p = 0.399). Similarly, the Waltz was associated with less Enjoyment than the Rock composition (t(444) = -3.76, p = 0.0011) with a trend for less Enjoyment compared to the Salsa composition (t(444) = -2.35, p = 0.088). There were no differences in Enjoyment for the Salsa and Rock compositions (t(444) = 1.41, p = 0.495). Pairwise comparisons for music playing conditions, averaged over the levels of song and corrected for multiple tests, revealed increased Enjoyment for the Improvise as compared to the Beat condition, for both Live (t(444) = 9.29, p < 0.00001) and Recorded (t(444) = 3.90, p = 0.0007) accompaniment conditions. Additionally, Enjoyment increased for the Live as compared to the Recorded condition, for both Improvise (t(444) = 8.17, p < 0.00001) and Beat (t(444) = -2.78, p = 0.03) music playing conditions. Finally, the combination of Improvise with Live conditions produced the highest levels of Enjoyment compared to the other condition combinations, including Improvise with Recorded (t(444) = 8.17, p < 0.00001), Beat with Live (t(444) = -9.29, p<0.00001), and Beat with Recorded (t(444) = -12.07, p<0.00001) condition combinations.

**Figure 3:**
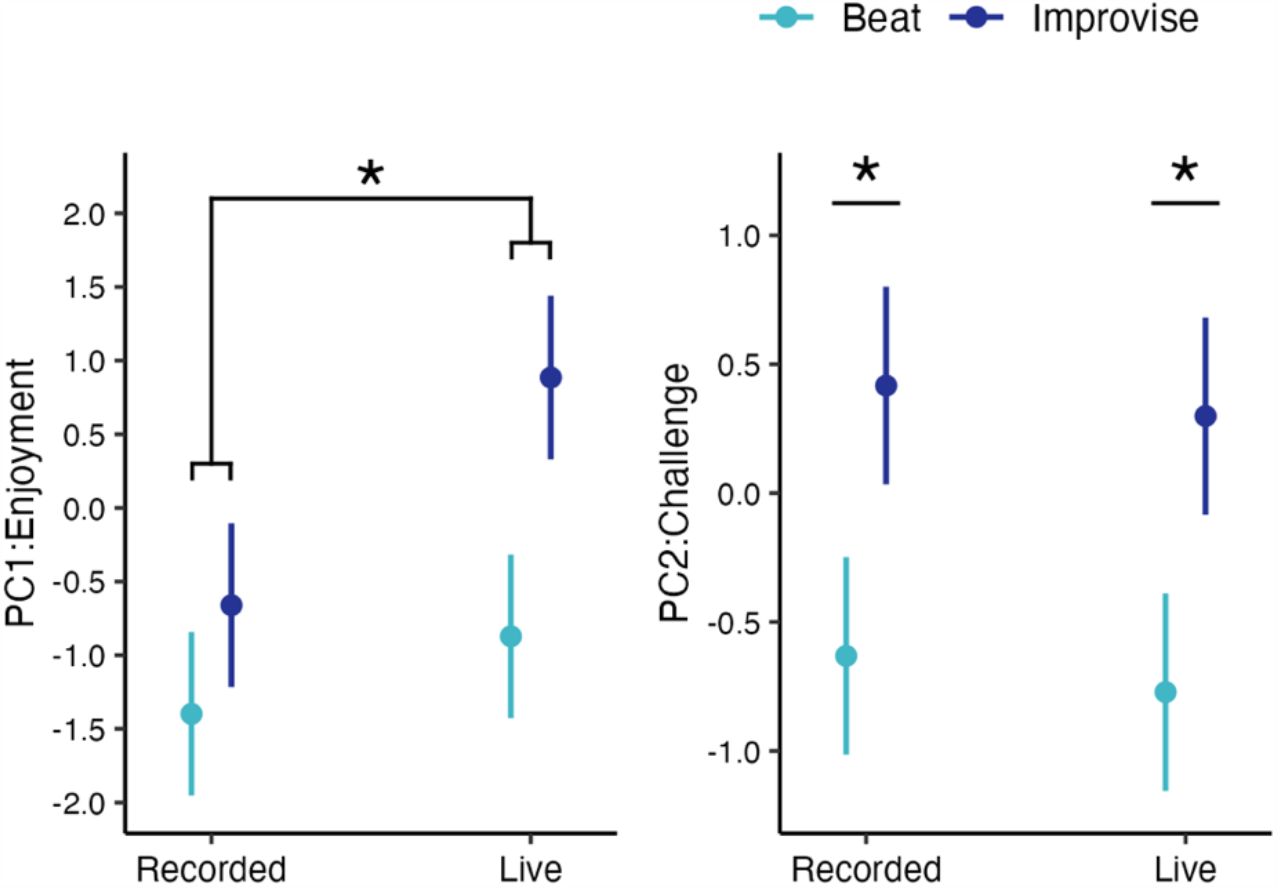
Behavioral Correlates of Music Playing Experience. There was a significant interaction between music playing condition and music accompaniment condition on Enjoyment, such that the combination of Improvise and Live conditions produced the greatest Enjoyment compared to the other three condition combinations. There was a main effect of music playing condition on Challenge, such that Challenge was greater during the Improvise as compared to the Beat condition. * p < 0.05 after correction for multiple tests.

With Challenge as the outcome (see Figure 3), there was a significant main effect for music playing condition (χ^2^ (1) = 50.12, p < 0.00001) and song (χ^2^ (3) = 35.42, p < 0.00001). There was no significant interaction between music playing condition and accompaniment condition (χ^2^ (1) = 0.011, p = 0.916) or a significant main effect of accompaniment (χ^2^ (1) = 0.897, p = 0.344), music training (χ^2^ (1) = 0.639, p = 0.424), or sensitivity to musical reward (χ^2^ (1) = 1.16, p = 0.281). Pairwise comparisons for song, corrected for multiple tests, revealed greater Challenge for the Salsa composition as compared to the March (t(444) = 4.72, p = 0.00002), Waltz (t(444) = 5.39, p = 0.0000007), and Rock (t(444) = 4.14, p = 0.00024) compositions. Pairwise comparisons for music playing condition, averaged over the levels of song and corrected for multiple tests, revealed higher levels of Challenge for the Improvise as compared to Beat condition, for both Live (t(444) = -7.23, p < 0.00001) and Recorded (t(444) = -7.08, p < 0.00001) accompaniment conditions.

### Motor Response to Music Conditions

For total acceleration (see Figure 4), there were no main effects for music playing condition (χ^2^(1) = 0.766, p = 0.382) or accompaniment condition (χ^2^(1) = 0.0005, p = 0.982), music training (χ^2^(1) = 0.727, p = 0.394), or sensitivity to musical reward (χ^2^(1) = 1.16, p = 0.281), although there was a significant main effect for song (χ^2^ (3) = 23.99, p = 0.00003). Importantly, there was a significant interaction between music playing condition and accompaniment condition on total acceleration (Figure 4; χ^2^ (1) = 7.54, p = 0.006). Pairwise comparisons for song, controlling for multiple tests, revealed lower total acceleration for the March as compared to the Rock (t(432) = -3.98, p = 0.0005), Salsa (t(432) = -4.41, p = 0.0001, and Waltz (t(432) = - 3.18, p = 0.0087) compositions. There were no significant differences in total acceleration between the Rock and Salsa (t(432) = -0.376, p = 0.982) or Waltz compositions (t(432) = 750, p = 0.877), or between the Salsa and Waltz compositions (t(432) = 1.125, p = 0.674). Pairwise comparisons for music playing condition, averaged over the levels of song and controlling for multiple tests, revealed that the combination of the Improvise with Live conditions produced greater total acceleration than the other three condition combinations, including Improvise with Recorded (t(432) = 3.90, p = 0.006), Beat with Live (t(432) = 4.76, p = 0.00002), and Beat with Recorded (t(432) = 4.78, p = 0.00001) condition combinations.

**Figure 4:**
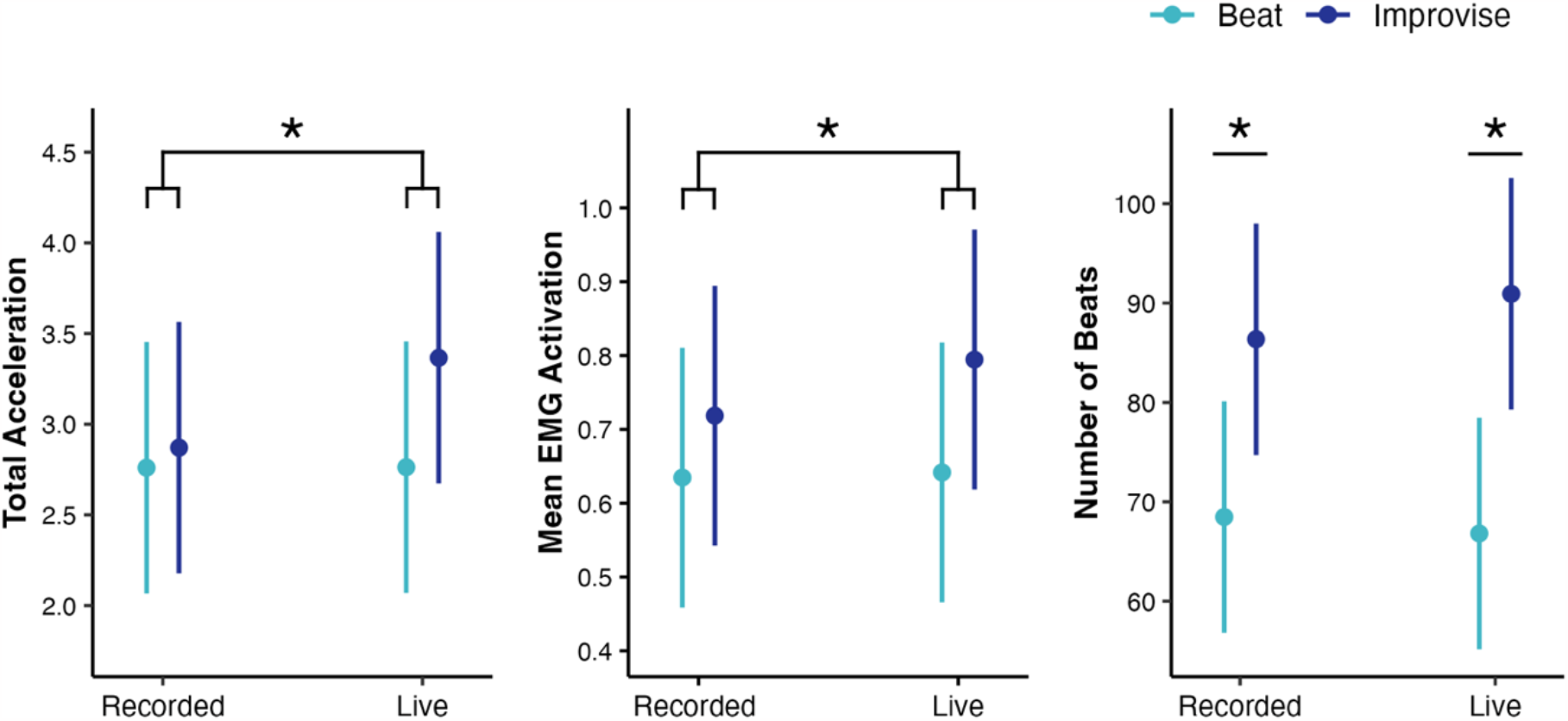
Motor Response to Music Conditions. There was a significant interaction between music playing condition and accompaniment condition on total acceleration and mean EMG activation, such that the combination of Improvisation and Live conditions produced the greatest motor response. There was a main effect of music playing condition on the number of beats played, such that a greater number of beats were produced during the Improvise as compared to Beat condition. * p < 0.05 after correction for multiple tests.

For mean EMG activation (see Figure 4), there were main effects of music playing condition (χ^2^(1) = 18.11, p = 0.00002) and song (χ^2^(3) = 82.60, p < 0.00001). There were no significant main effects of accompaniment condition (χ^2^(1) = 0.13, p = 0.720), musical training (χ^2^(1) = 0.30, p = 0.581), or sensitivity to musical reward (χ^2^(1) = 0.05, p = 0.816). Importantly, there was also an interaction effect between music playing condition and accompaniment condition on mean EMG activation (Figure 4; χ^2^ (1) = 6.12, p = 0.013). Pairwise comparisons for song, controlling for multiple tests, revealed less EMG activation for the March as compared to the Rock (t(372.08) = -6.16, p < 0.00001) and Salsa (t(372.00) = -6.35, p < 0.00001) compositions, but not the Waltz (t(372.12) = 0.461, p = 0.967). Similarly, there was lower EMG activation for the Waltz as compared to the Rock (t(372.04) = -6.50, p < 0.00001) and Salsa (t(372.11) = - 6.63, p < 0.00001) compositions. There were no significant differences in EMG activation between the Rock and Salsa compositions (t(372.08) = 0.097, p = 0.9997). Pairwise comparisons of music playing conditions, averaged over the levels of song and controlling for multiple comparisons, showed greater mean EMG activation for the Improvise as compared to Beat conditions for both Live (t(372) = -7.76, p < 0.00001) and Recorded (t(372) = 4.26, p = 0.0002) accompaniment conditions. In addition, the combination of the Improvise with Live conditions produced greater mean EMG activation compared to the other three condition combinations, including Improvise with Recorded (t(372) = 3.86, p = 0.0008), and Beat with Live (t(372) = 7.76, p < 0.00001), and Beat with Recorded (t(372) = 8.11, p < 0.00001) condition combinations.

For the number of beats played (see Figure 4), there were main effects of music playing condition (χ^2^ (1) = 23.62, p < 0.00001) and song (χ^2^ (3) = 61.13, p = 0.00001). There were no significant main effects of accompaniment condition (χ^2^ (1) = 0.20, p = 0.652), musical training (χ^2^(1) = 0.01, p = 0.909), nor a significant interaction between music condition and accompaniment condition (χ^2^ (1) = 1.43, p = 0.232). There was a trend towards a main effect of sensitivity to musical reward on the number of beats played (χ^2^(1) = 3.64, p = 0.056). Pairwise comparisons for song, controlling for multiple tests, revealed fewer number of beats played for the March, as compared to the Rock (t(444) = -5.47, p = 0.0000005), Salsa (t(444) = -7.39, p < 0.0000001), and Waltz (t(444) = -5.53, p = 0.0000003) compositions. There were no significant differences in the number of beats played between the Rock and Salsa (t(444) = -1.92, p = 0.220) or Waltz (t(444) = -0.066, p = 0.9998) compositions, or between the Salsa and Waltz compositions (t(444) = 1.86, p = 0.249). Pairwise comparisons in music playing condition, averaged over the level of song and controlling for multiple tests, revealed a greater number of beats played during the Improvise as compared to Beat conditions, for both the Live (t(444) = 4.86, p < 0.00001) and Recorded (t(444) = 6.55, p < 0.00001) accompaniment conditions.

### Autonomic Response to Music Conditions

For tonic EDA (see Figure 5), there was a main effect of music playing condition (χ^2^ (1) = 5.82, p = 0.016), while controlling for the effects of movement by including total acceleration in the model. There was also a significant main effect of total acceleration (χ^2^ (1) = 5.01, p = 0.026) and song (χ^2^ (3) = 8.33, p = 0.040). There were no main effects of accompaniment condition (χ^2^(1) = 0.05, p = 0.830), musical training (χ^2^(1) = 0.53, p = 0.467), or sensitivity to musical reward (χ^2^(1) = 0.03, p = 0.874), or any significant interaction between music playing and accompaniment conditions (χ^2^(1) = 0.66, p = 0.418). Pairwise comparisons for song, controlling for multiple testing, revealed lower tonic EDA for the March as compared to the Salsa composition (t(361.64) = -2.64, p = 0.043), but not the Rock (t(363.44) = -0.93, p = 0.788) or Waltz (t(361.42) = -2.04, p = 0.174) compositions. There were no significant differences between the Rock and Salsa (t(357.14) = -1.70, p = 0.325) or Waltz (t(357.02) = -1.12, p = 0.675) compositions, or between the Salsa and Waltz compositions (t(357.77) = 0.54, p = 0.948). Pairwise comparisons for music playing conditions, averaged over the levels of song and controlling for multiple testing, revealed greater tonic EDA, indicating higher autonomic arousal, for the Improvise as compared Beat condition for both Live (t(361.18) = 3.51, p = 0.0005) and Recorded (t(356.15) = 2.41, p = 0.016) accompaniment conditions.

**Figure 5:**
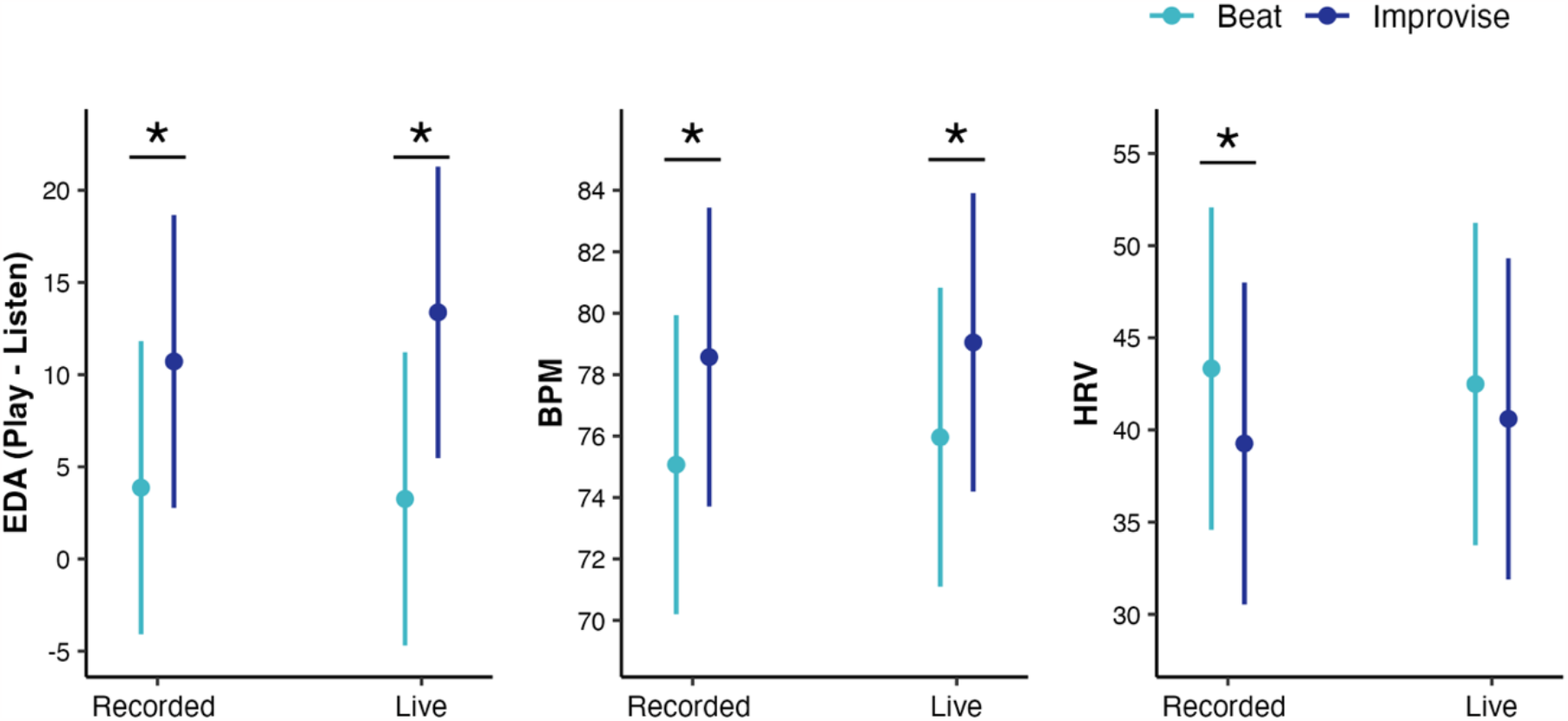
Autonomic Response to Music Conditions. There was a main effect of music playing condition on autonomic response, such that the Improvise condition led to increased autonomic arousal as measured by increased tonic EDA and BPM, and parasympathetic withdrawal as measured by decreased HRV, as compared to the Beat condition. Pairwise comparisons revealed higher EDA and BPM for the Improvise as compared to Beat condition during both Recorded and Live accompaniment conditions. For HRV, the difference between the Improvise and Beat conditions was only significant in the Recorded condition. * p < 0.05 after correction for multiple tests.

For BPM (see Figure 5), there was a main effect of music playing condition (χ^2^ (1) = 21.34, p < 0.00001), while controlling for the effects of movement by including total acceleration in the model. There were also significant main effects of total acceleration (χ^2^ (1) = 13.45, p = 0.0002) and musical training (χ^2^ (1) = 4.35, p = 0.037). There were no main effects of accompaniment condition (χ^2^(1) = 1.39, p = 0.238), song (χ^2^ (3) = 7.48, p = 0.058), or sensitivity to musical reward (χ^2^(1) = 0.42, p = 0.519), or any significant interaction between music playing condition and accompaniment condition (χ^2^(1) = 0.15, p = 0.701). Pairwise comparisons for music playing condition, averaged over the level of song and controlling for multiple testing, revealed greater BPM, indicating higher autonomic arousal, for the Improvise as compared to Beat conditions for both Live (t(398.65) = 3.95, p < 0.00001) and Recorded (t(397.07) = 4.62, p < 0.00001) accompaniment conditions.

For HRV (see Figure 5), there was a main effect of music playing condition (χ^2^ (1) = 4.66, p = 0.041), while controlling for the effects of movement by including total acceleration in the model. There was also a significant main effect of total acceleration (χ^2^ (1) = 10.16, p = 0.0014). There were no main effects of accompaniment condition (χ^2^(1) = 0.20, p = 0.655), song (χ^2^(3) = 3.91, p = 0.271), musical training (χ^2^(1) = 3.67, p = 0.055), or sensitivity to musical reward (χ^2^(1) = 0.002, p = 0.961), or any significant interaction between music playing and accompaniment conditions (χ^2^(1) = 0.66, p = 0.418). Pairwise comparisons for music playing condition, averaged over the levels of song and controlling for multiple testing, revealed lower HRV, indicating parasympathetic withdrawal (Mackersie and Calderon-Moultrie, 2016), for the Improvise compared to Beat conditions for the Recorded accompaniment condition (t(397.14) = 2.16, p = 0.031). Although HRV was also lower for the Improvise compared to Beat conditions for the Live accompaniment condition, this difference was not significant (t(400.12) = 0.976, p = 0.330).

### Association Between Enjoyment and Motor Responses

The association between Enjoyment and each motor response (total acceleration, mean EMG activation, and number of beats played) was evaluated by adding the motor response to the linear mixed model predicting Enjoyment. The main effect of motor response was considered, as well as the three-way interaction between motor response, music playing condition, and accompaniment condition, the main effect for control variables for song, musical training, and sensitivity to musical reward, and a random intercept for subject. To account for multiple comparisons, significance levels with and without Bonferroni correction are reported for the association between Enjoyment and the three motor responses.

There was no significant main effect of total acceleration on Enjoyment (χ^2^ (1) = 0.04, p = 0.841) and no significant triple interaction between total acceleration, music playing condition, and accompaniment condition on Enjoyment (χ^2^ (1) = 0.0004, p = 0.983). However, there was a significant interaction effect between total acceleration and music playing condition on Enjoyment (Figure 6; χ^2^ (1) = 12.15, p = 0.0005). This effect remained significant after Bonferroni correction for multiple comparisons (p = 0.0015). There was also a significant main effect of song (χ^2^(3) = 28.71, p = 0.000003) and sensitivity to musical reward (χ^2^(1) = 3.92, p = 0.048). . There were no significant interaction effects for total acceleration and accompaniment condition (χ^2^ (1) = 0.098, p = 0.755) or music playing condition and accompaniment condition (χ^2^ (1) = 3.05, p = 0.081, and no main effects of musical training (χ^2^ (1) = 0.533, p = 0.465), accompaniment condition (χ^2^ (1) = 3.46, p = 0.063), or music playing condition (χ^2^ (1) = 3.34, p = 0.841). Notably, the previously significant interaction between music playing condition and accompaniment condition and the main effect of music playing condition on Enjoyment became non-significant after including total acceleration in the model. Pairwise comparisons of song on Enjoyment have been reported above and maintained the same relationships. Simple effects for the association between total acceleration and revealed a positive relationship between total acceleration and Enjoyment during the Improvise condition, using the 95% confidence interval to indicate significance (estimated slope = 0.302; 95% CI = 0.166, 0.438), but not for the Beat condition (estimated slope = -0.003; 95% CI = -0.153, 0.148). The slopes describing the relationship between Enjoyment and total acceleration were significantly different between the Improvise and Beat conditions (estimate = 0.305, t(453) = 4.60, p < 0.0001).

**Figure 6:**
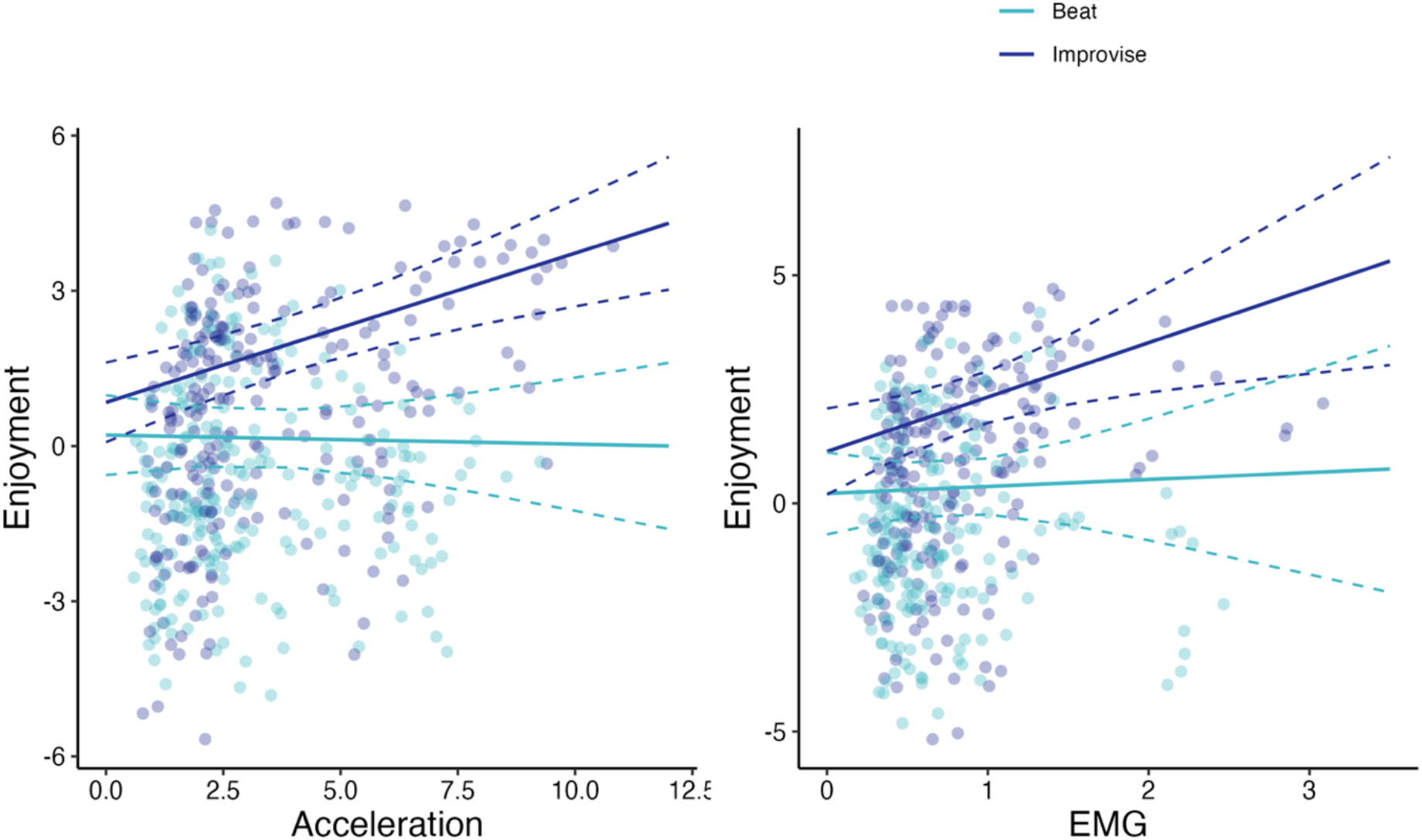
Relationship between reward and motor outputs. There was a significant interaction between total acceleration and music playing condition on Enjoyment, such that Enjoyment was positively associated with total acceleration for the Improvise condition but not the Beat condition. Similarly, there was a significant interaction between mean EMG activation and music playing condition on Enjoyment, such that Enjoyment was positively associated with mean EMG activation for the Improvise condition but not the Beat condition.

There was no significant main effect of mean EMG activation on Enjoyment (χ^2^ (1) = 0.10, p = 0.753) and no significant triple interaction effect between mean EMG activation, music playing condition, and accompaniment condition on Enjoyment (χ^2^ (1) = 0.30, p = 0.586). However, there was a significant interaction effect between mean EMG activation and music playing condition on Enjoyment (Figure 6; χ^2^(1) = 5.55, p = 0.0184), while the interaction between music playing condition and accompaniment condition approached significance (χ^2^ (1) = 3.78, p = 0.052). After Bonferroni correction, the interaction between mean EMG activation and music playing condition on Enjoyment approached significance (p = 0.0552). There was also a significant main effect of music playing condition (χ^2^ (1) = 5.54, p = 0.019), song (χ^2^(3) = 19.90, p = 0.0002), and sensitivity to musical reward (χ^2^(1) = 8.89, p = 0.003). There was no significant interaction effect for mean EMG activation and accompaniment condition (χ^2^ (1) = 0.07, p = 0.786) and no main effects of musical training (χ^2^ (1) = 0.63, p = 0.426) or accompaniment condition (χ^2^ (1) = 2.40, p = 0.122). Pairwise comparisons of song on Enjoyment have been reported above and maintained the same relationships. Simple effects for the association between mean EMG and Enjoyment revealed a positive relationship between mean EMG activation and Enjoyment during the Improvise condition, using the 95% confidence interval to indicate significance (estimated slope = 1.300; 95% CI = 0.483, 2.117) but not the Beat condition (estimated slope = 0.091; 95% CI = -0.784, 0.967). The slopes describing the relationship between Enjoyment and mean EMG activation were significantly different between Improvise and Beat conditions (estimate = 1.21, t(391) = 3.81, p = 0.0002).

There was no significant main effect of beats played on Enjoyment (χ^2^ (1) = 0.84, p = 0.360) and no significant triple interaction effect between beats played, music playing condition, and accompaniment condition on Enjoyment (χ^2^(1) = 0.0065, p = 0.936). The interaction effect between beats played and music playing condition on Enjoyment approached significance (χ^2^ (1) = 3.83, p = 0.0503), but was not significant after Bonferroni correction (p = 0.151). There was also a significant main effect of song (χ^2^(3) = 26.08, p = 0.000009), and a trend towards a main effect for sensitivity to musical reward (χ^2^(1) = 3.05, p = 0.081). There were no significant interaction effects for beats played and accompaniment condition (χ^2^ (1) = 0.204, p = 0.651) or music playing condition and accompaniment condition (χ^2^ (1) = 2.04, p = 0.154), and no main effects of musical training (χ^2^ (1) = 0.247, p = 0.619), accompaniment condition (χ^2^ (1) = 2.56, p = 0.109), or music playing condition (χ^2^ (1) = 2.20, p = 0.138). Notably, the previously significant interaction between music playing condition and accompaniment condition and the main effect of music playing condition on Enjoyment became non-significant after including beats played in the model. Pairwise comparisons of song on Enjoyment have been reported above and maintained the same relationships. Simple effects for the association between beats played and Enjoyment revealed a positive relationship between beats played and Enjoyment during the Improvise condition, using the 95% confidence interval to indicate significance (estimated slope = 0.0133; 95% CI = 0.0078, 0.0189), but not the Beat condition (estimated slope = 0.0046; 95% CI = -0.0012, 0.0104). The slopes describing the relationship between Enjoyment and beats played were significantly different between the Improvise and Beat conditions (estimate = 0.0087, t(496) = 2.53, p = 0.0118).

## Discussion

Here we show that improvisation and live accompaniment increase enjoyment and motor performance during a music playing task . The combination of improvisation and live accompaniment led to increased acceleration of hand movements while playing the drum, coupled with increased amplitude of muscle contractions in wrist flexor and extensor muscles. Importantly, increased movement was associated with increased reported Enjoyment only when improvising. These findings suggest a mechanism by which combined improvisation and live accompaniment increase Enjoyment and motivate increased motor activity. We also show that improvisation leads to increased autonomic arousal, as measured by increased tonic EDA and BPM and decreased HRV. Together, these findings have implications for enhancing reward, engagement, and motor performance during music playing tasks that support exercise and physical rehabilitation.

In our experimental design, we did not provide any instructions to participants regarding how much they should move while playing music in any condition. The increased acceleration and muscle activation during conditions with combined improvisation and live accompaniment might thus reflect participants’ intrinsic motivation to move. Supporting this mechanism, participants’ self-reports of pleasure, motivation, momentum, and activation loaded onto a principal component that we termed “Enjoyment” which increased for conditions with improvisation and live accompaniment as compared to the other conditions. Further, the association between Enjoyment and motor response was moderated by the music playing condition, such that increased Enjoyment was associated with increased total acceleration of hand movements while improvising, but not while maintaining the beat. The same moderation effect approached significance for mean EMG activation, with simple effects demonstrating that increased mean EMG activation was significantly associated with increased Enjoyment while improvising, but not while maintaining the beat. This relationship between Enjoyment and movement is supported by previous research demonstrating that reward enhances motor performance and learning (Terry et al., 2020) and contributes to motor recovery post-stroke (Grau-Sanchez et al., 2018). An association between enhanced music reward and improved motor learning has also been identified in a music playing task involving playing a learned melody (Bianco et al., 2019). In our task, the flexibility of choice provided during improvisation may have played a key role in enhancing reward and movement. Autonomy of choosing what to do has been shown to enhance reward, motor performance, and learning (Wulf et al., 2014; Lewthwaite et al., 2015; Murayama et al., 2015; Chua et al., 2018; An et al., 2020). However, it is also possible that the limited variation in movement while maintaining the beat inhibited reward-enhanced movement for this condition. Together, our findings demonstrate that improvisation and live accompaniment may be used to facilitate increased Enjoyment and movement during music playing, and that the association between reward and movement is greater during improvisation.

The music playing condition also produced a main effect on average muscle activation, the number of beats played, and Enjoyment, with the Improvise condition associated with greater motor response and Enjoyment as compared to the Beat condition. This suggests that improvisation, regardless of live or recorded accompaniment, may increase reward and enhance motor performance. Participants chose their own beat, and therefore set their own pace when improvising and maintaining the beat; however, they increased their frequency of movement while improvising, as demonstrated by the increase in the number of beats played. This increase in frequency during improvisation may be partially due to the freedom from constraints in movement frequency, which necessarily limited the number of beats that could be played during the Beat condition. However, the EMG analysis addressed this limitation by averaging peak EMG activation for each movement, in this way measuring movement intensity independent of movement frequency. This analysis revealed increased muscle activation for the Improvise condition as compared to the Beat condition, supporting the link between improvisation and enhanced motor performance. Together, these findings suggest an important role of improvisation for enhancing reward and motor performance, with mechanisms that may relate to increased autonomy and agency (Wulf et al., 2014; Lewthwaite et al., 2015; Murayama et al., 2015; Chua et al., 2018; An et al., 2020).

The accompaniment condition produced a main effect on Enjoyment, with live accompaniment producing the greater Enjoyment compared to recorded accompaniment. However, there were no main effects of the accompaniment condition on motor or autonomic responses. This suggests that the benefits of live accompaniment are best realized in combination with improvisation, as the interaction between live accompaniment and improvisation led to the greatest levels of Enjoyment and enhanced motor performance. Social cooperation has demonstrated benefits for improving intrinsic motivation and motor learning (Kaefer and Chiviacowsky, 2022), and the present findings suggest that the combined effects of live accompaniment and improvisation supported a cooperative effort to coordinate drum and piano playing. Importantly, the piano accompaniment for this study was provided by a music therapist with specialized training in clinical improvisation, which emphasizes responsiveness to participant playing to facilitate a shared music playing experience (Nordoff et al., 2007). Together, these results suggest that live accompaniment, on its own, does not enhance motor performance, but that live accompaniment can enhance the benefits of improvisation for motor performance through a mechanism of intrinsic reward that might be related to social cooperation.

We also demonstrated that improvisation during music-playing increased autonomic arousal, as measured via tonic EDA, BPM, and HRV, as well as the experience of Challenge. Importantly, the association between improvisation and autonomic arousal remained significant after controlling for movement, suggesting that arousal relates to mechanisms beyond physical exertion (Sara and Bouret, 2012). Given the association between improvisation and Challenge, it may be that the increased autonomic arousal during improvisation relates to cognitive control. Supporting this potential mechanism, rhythmic improvisation has previously been linked to increased activation in the dorsal anterior cingulate cortex (dACC), when compared to repeating a rhythm (de Aquino et al., 2019), and previous research links dACC activation to cognitive control (Niendam et al., 2012) and evaluation of creative tasks (Ellamil et al., 2012). However, music improvisation has also been linked to activation in the default mode network and deactivation of the executive network, with the extent of executive network deactivation shown to vary with the level of expertise in music improvisation (Pinho et al., 2014; Beaty, 2015).

These findings highlight the complexity of improvisation as a creative task that involves both idea generation and evaluation (Ellamil et al., 2012; Loui, 2018). Our findings suggest that cognitive challenge played a role in rhythmic improvisation for our participants, an effect that remains significant while controlling for the level of musical expertise.

Finally, our analyses demonstrate that the song composition significantly modulated Enjoyment, Challenge, motor response, and autonomic response during music playing. Familiarity with song genres and the amount of musical groove may have played a role in modulating responses to each song. Although musical preference is thought to vary between individuals, the effects of song composition in this study suggests future research should examine how to manipulate song characteristics to enhance motor and affective outcomes during music playing (Matthews et al., 2020; Park et al., 2019; Periera et al., 2011; Witek et al., 2014).

Together, our findings suggest two distinct mechanisms influencing response to music-playing tasks. First, Enjoyment during music-playing is facilitated by the combination of improvisation and live accompaniment. During improvisation, Enjoyment is associated with increased acceleration of hand movements and approaches significant association with muscle activation. Second, improvisation increases Challenge and autonomic arousal while controlling for motor output, suggesting increased attentional processes when generating musical ideas. These findings have important implications for increasing reward and attention during music-playing tasks, which may promote engagement, dose of movement, and improved outcomes during exercise and physical rehabilitation. Understanding the link between music reward and movement is essential for improving our general understanding of the intricate relationship between motor and reward-related regions and their neural substrates (Aoki et al., 2019). Future studies should use neuroimaging methods to investigate functional connectivity between reward and sensorimotor regions during music playing, and to better understand the relationship between improvisation, cognitive challenge, and executive function. Additionally, studies investigating these mechanisms in clinical populations have potential to inform the development of music playing tasks to optimize motor rehabilitation (Braun Janzen et al., 2022).

## EXTENDED DATA

**Figure 1.1:**
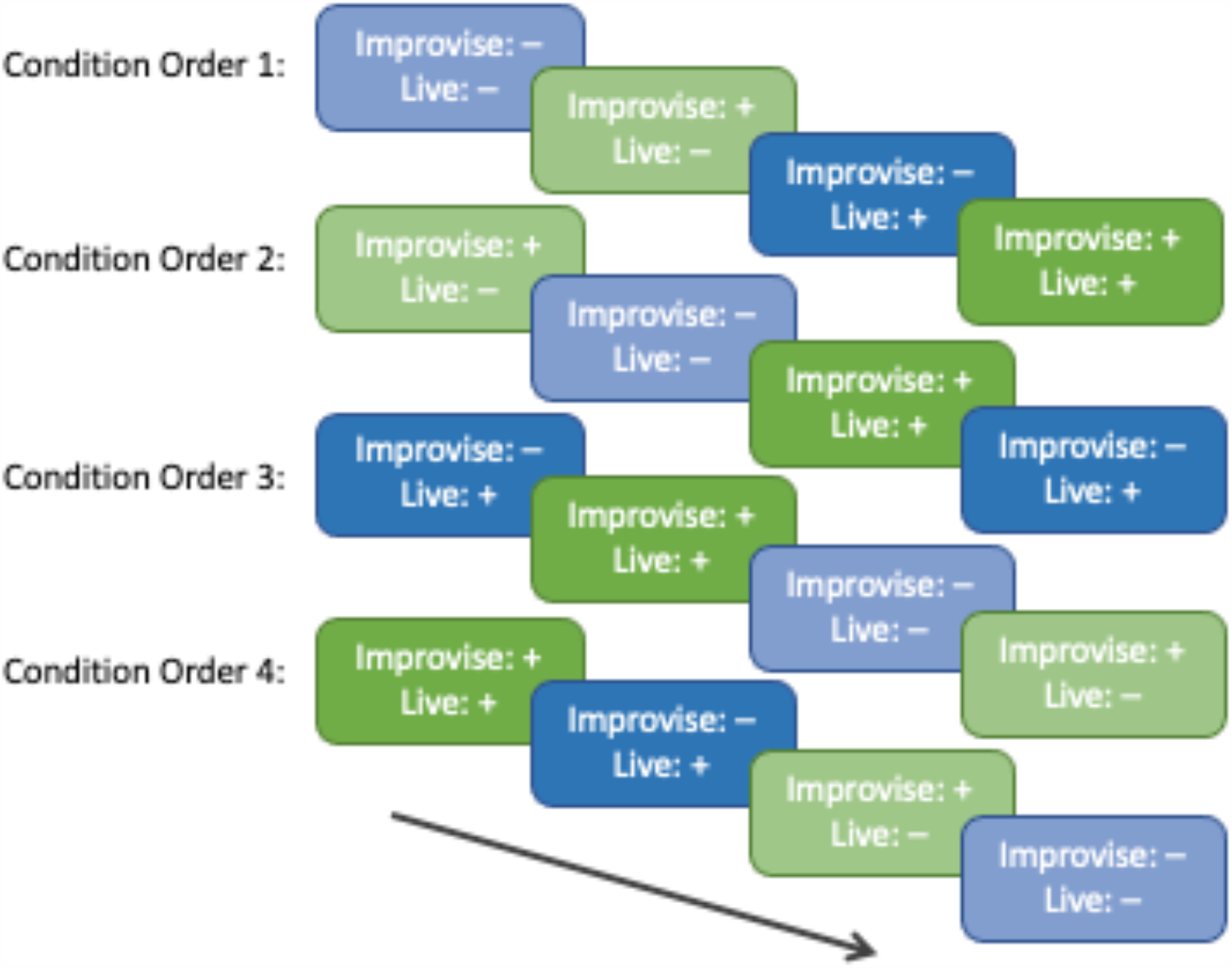
Condition Orders Participants were randomized to one of four condition orders. Each condition order alternated between music playing conditions (Improvise vs. Beat), with half of the condition orders starting with the Improvise condition. Accompaniment conditions (Live vs. Recorded) were completed consecutively, such that participants completed both music playing conditions with one accompaniment condition and then repeated both music playing conditions with the other accompaniment condition. The same condition order was repeated for each of the four songs played by each participant.

