## Extended Data for "Improvisation and Live Accompaniment Increase Motor Response and Reward During a Music Playing Task"

**Figure 1.1**: Condition Orders

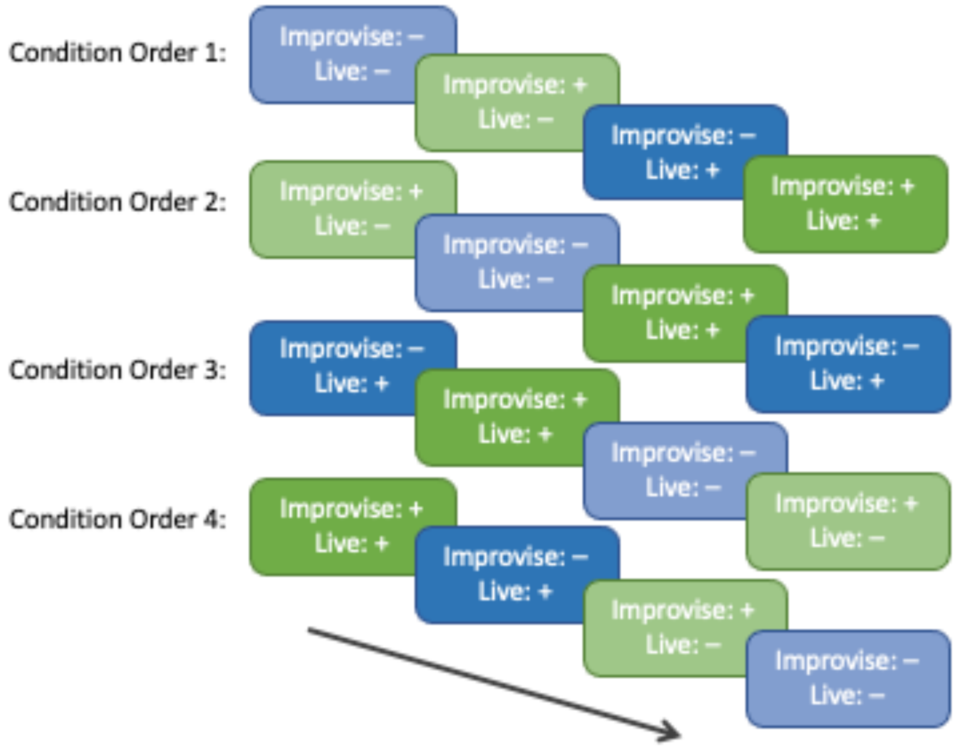


Participants were randomized to one of four condition orders. Each condition order alternated between music playing conditions (Improvise vs. Beat), with half of the condition orders starting with the Improvise condition. Accompaniment conditions (Live vs. Recorded) were completed consecutively, such that participants completed both music playing conditions with one accompaniment condition and then repeated both music playing conditions with the other accompaniment condition. The same condition order was repeated for each of the four songs played by each participant.
